# Quantifying epigenetic reprogramming and cellular plasticity in preneoplastic lesions reveals dedifferentiation as a key precancer hallmark

**DOI:** 10.64898/2026.10.25.761640

**Authors:** Jason Tham Han Kiat, Andrew E. Teschendorff

**Affiliations:** Shanghai Institute of Nutrition and Health, University of Chinese Academy of Sciences, Chinese Academy of Sciences, Shanghai 200031, China

**Keywords:** single cell RNA-Seq, cancer risk, cancer stem cell, transcription factor, regulatory network, metaplasia

## Abstract

Epigenetic reprogramming and plasticity are two cancer hallmarks that increase the risk of carcinogenic transformation, yet quantifying these properties remains a fundamental challenge. Here we develop and validate a general pan-tissue computational framework called PLASTIC for quantifying cellular plasticity and epigenetic reprogramming from single-cell RNA-seq data. PLASTIC allows construction of Waddington-like landscapes allowing identification of high cancer risk states. When applied to a pan-preneoplastic tissue setting, PLASTIC reveals how the level of epigenetic reprogramming of precancerous cells correlates with cellular plasticity and cancer-risk, and that the high cancer risk state is characterized by increased dedifferentiation, even when compared to its fetal cell counterpart. Furthermore, PLASTIC predicts the emergence of metaplasia in normal lung cells from smokers, identifying a high cancer risk signature that we validate in independent data. In summary, PLASTIC can help quantify cellular plasticity and dedifferentiation of single cells in any tissue-type, opening up new strategies for cancer risk prediction and prevention.

## Introduction

The most recent enumeration of cancer hallmarks lists unlocking phenotypic plasticity and non-mutational epigenetic reprogramming as two key cellular properties that enable, or facilitate, the acquisition of other more established hallmarks such as epithelial-mesenchymal transformation (EMT), invasion, or evasion of the immune system [1–3]. Recent work supports the view that increased plasticity and epigenetic heterogeneity are key drivers of cancer progression [4–15]. Increased epigenetic and phenotypic plasticity may however also play a key role in the earliest stages of cancer development, predisposing normal cells to carcinogenic transformation [16–18]. Supporting this further is the observation that cancer is often preceded by various forms of lesions that display dedifferentiation, blocked differentiation or trans-differentiation [1, 2], collectively referred to as dysplasia or metaplasia. However, the precise relationship between increased cellular plasticity and these various forms of aberrant epigenetic reprogramming remains unclear. Accentuating this challenge, we currently also have a limited understanding as to which particular preneoplastic cells undergo carcinogenic transformation. A long-held view is that cancer arises from an aberrant dedifferentiated stem-like state [19–21] [22, 23] [24–28]. This cancer stem cell (CSC) hypothesis however has been questioned by studies demonstrating how in certain cancer-types, the founding cancer cell may arise from a differentiated cell that undergoes aberrant dedifferentiation [29]. Nevertheless, regardless of whether the founder cell is a stem-cell or a terminally differentiated cell that has undergone dedifferentiation, in both cases the founder cell is marked by dedifferentiation. Thus, put together, it would be important to develop strategies that can directly quantify the level of plasticity and dedifferentiation, and to do so within an unbiased systems-biological pan-tissue framework [30–33]. Whilst the advent and consolidation of single-cell technologies like scRNA-Seq and snRNA-Seq [34–37] offer the unprecedent opportunity to develop such strategies [38–40], a considerable challenge remains in how to computationally estimate cellular plasticity and dedifferentiation from what is noisy single-cell omic data.

Here we propose and validate a systems biological pan-tissue computational framework, called PLASTIC **(P**redictive **L**andscape **A**nalysis of **S**ingle-cell **T**ranscriptomes for **I**dentifying **C**ancer), to help quantify cellular plasticity and epigenetic reprogramming of premalignant lesions from scRNA-Seq or snRNA-Seq data. We use PLASTIC to quantify and delineate the cellular heterogeneity of metaplasia and other precancerous lesions at single-cell resolution in the context of progression to three cancer-types: intestinal-type gastric cancer, lung squamous cell carcinoma and colorectal adenocarcinoma. Using PLASTIC, we build one of the first examples of a Waddington landscape of pre-cancer states, illustrated here for the main cellular states found in intestinal metaplasia, and predicting that intestinal-type gastric cancer emerges from an abnormally transformed highly dedifferentiated and plastic goblet progenitor cell. In the case of normal lung basal cells from smokers, PLASTIC correctly predicts transformation into squamous metaplasia and identifies a high cancer risk state that we successfully validate in an independent multi-stage dataset of lung cancer progression. Overall, our data points towards the high cancer risk state being defined by common precancer hallmarks, that include a state of dedifferentiation characterized by loss of a tissue-specific fetal gene expression program, loss of cell-adhesion and an increased cellular plasticity marked by a conserved and cell-proliferation independent ribosomal gene expression program.

## Results

### PLASTIC: a Waddington framework to quantify cellular plasticity and epigenetic reprogramming

Since lineage specific transcription factors (TFs) orchestrate epigenetic reprogramming in normal development [41–43], measuring their activity in the cell-fate commitment and differentiation process offers a principled way to quantify epigenetic reprogramming generally. To identify lineage-specific TFs we use the SCIRA algorithm [44, 45], a machine learning method that is applied to the large multi-tissue RNA-Seq GTEX dataset [46], whilst adjusting for immune-cell infiltration, to infer a tissue-specific TF-differentiation network, comprising lineage-specific TFs and their regulons (**Methods, Fig.1a**). The genes making up these regulons are enriched for TF-binding targets but also include many indirect downstream targets that faithfully represent upstream TF-binding activity [47, 48]. Expression levels of these regulon genes can be used to estimate a TF’s differentiation activity (TDA) (**Methods**) [44, 49, 50], which, as shown by us in the context of several tissue-types [44, 50], significantly outperforms the TF-expression level as a proxy of a TF’s differentiation function binding activity. Whilst in previous work we have extensively validated such tissue-specific TF-differentiation networks for the normal lung, kidney, liver, pancreas [44] and esophagus [45] (**Fig.1a**), here we applied SCIRA to derive corresponding networks for normal stomach and colon (**SI tables S1-S2**), in order to then explore epigenetic reprogramming in human metaplasia and dysplasia, focusing on 3 well-known precancerous lesions [51] (**Fig.1b**): (i) gastric intestinal metaplasia, which precedes intestinal-type gastric cancer [52], (ii) squamous metaplasia in the lung of smokers, which precedes lung squamous cell carcinoma [51], and (iii) adenomatous polyps, which precedes colorectal adenocarcinoma [53]. For validation purposes, we also consider intestinal metaplasia of the esophagus (Barrett’s Esophagus), which normally precedes esophageal adenocarcinoma [54]. Since GTEX profiled adult normal tissues, the inferred TFs should be implicated mostly in maintaining differentiation, i.e homeostasis of the corresponding adult tissue. However, we found that many of the inferred TFs are also critical for early development of the corresponding tissue-types. For instance, the stomach differentiation network contained 52 TFs, including many well-known gastric specific TFs [55], such as *FOXQ1* (a pit mucus cell differentiation marker), *XBP1* (a chief cell maturation marker), *SPDEF* (mucous cell marker), *BARX1* (gastric mesoderm) and *SOX9* (pylorus marker), with many of the other identified TFs (e.g. *SOX2, GATA4, GATA6, FOXA1, FOXA2, HNF1A*) playing more critical roles in the early development of the stomach endoderm (**SI table S1**) [56]. The colon differentiation network contained 56 TFs, including well-known colon specific factors like *CDX2, CDX1*, *KLF4* and *ATOH1,* and others that were shared with stomach (e.g. *FOXA1, FOXA2, KLF5, TRIM31*) (**SI table S2**) [57]. However, a number of inferred TFs may not be specific to the resident epithelial cells of the tissue, but rather to tissue-specific stromal cells (e.g. fibroblasts or endothelial cells), a limitation we address here by filtering out these TFs with scRNA-Seq atlas data of the Tabula Sapiens [58], resulting in 47 stomach-epithelial and 18 colon-epithelial specific TFs (**Fig.1c, SI fig.S1-S2, SI tables S1-S2**). Using the ChIP-Seq Atlas [59] and the DOROTHEA regulome database [60], we verified that the regulons of stomach and colon epithelial specific TFs are enriched for direct TF-binding targets (**SI fig.S3**). By using these regulons to estimate TDA, we can thus accurately delineate the differentiation state of any given cell, including cells from precancerous states like metaplasia or dysplasia, as well as cancer tissue, representing them in a two-dimensional diffusion-map space [61] (**Fig.1d**). This diffusion-map in TDA-space then offers a means to approximate a 3D Waddington-like landscape if integrated with a metric that can quantify the elevation in the landscape, representing either differentiation potential (for a physiologically normal state) or, more generally, cellular plasticity. To approximate cellular plasticity we use a measure of transcriptomic entropy called diffusion or signaling entropy [62, 63] (**Fig.1e**), which correlates with differentiation potential and drug resistance, two key manifestations of cellular plasticity, in the context of both bulk mRNA expression [64, 65] and single-cell RNA-Seq data [62, 63]. Specifically, we use a computationally efficient proxy to diffusion entropy called CCAT **(Fig.1e**), which has been extensively validated by us and others [66–68]. Integrating this measure of cellular plasticity with our 2D diffusion map representation of TDA-space then yields a Waddington-like landscape with attractor states representing normal differentiated, precancer and cancer-states, allowing identification of precancer-states that are proximal to cancer and thus at potentially higher cancer-risk (**Fig.1f**, **Methods**).

**Figure-1:**
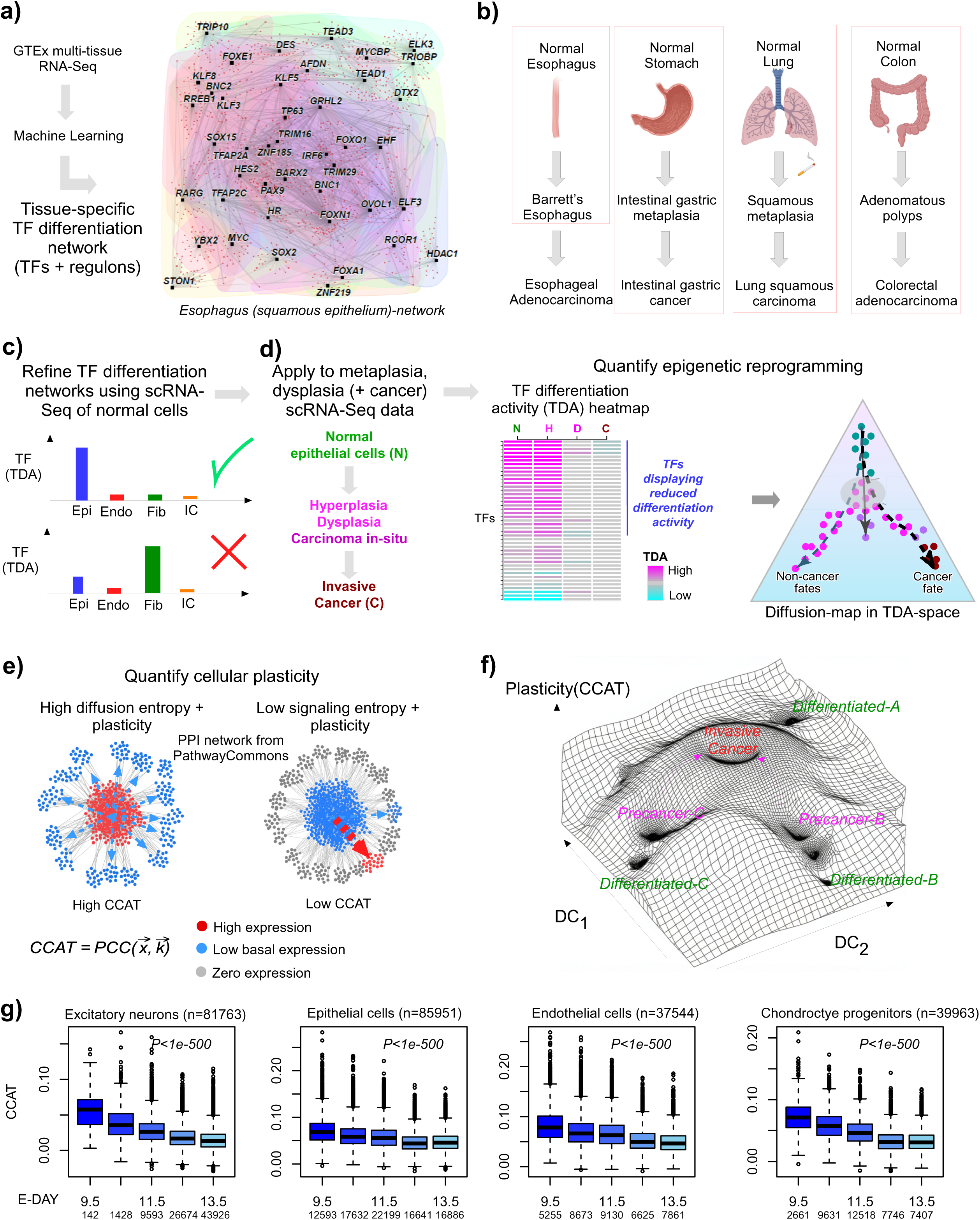
The PLASTIC computational framework for identifying high cancer-risk states. **a)** The first step involves the construction of a tissue-specific transcription factor (TF) differentiation network, which is inferred from the large multi-tissue GTEX RNA-Seq dataset using a machine learning algorithm called SCIRA, adjusting for immune-cell infiltration. This results in a set of tissue-specific TFs and associated regulons. As an example, the TF differentiation network for the normal esophagus is shown. **b)** In this work we analyze scRNA-seq data from precancerous lesions in the context of development of 4 cancer-types (enclosed in red boxes), as shown. This means, in step-a) we build TF-differentiation networks for esophagus (squamous epithelium), stomach-epithelium, lung-epithelium and colon-epithelium. **c)** The tissue-specific TF differentiation networks built in step a) are then refined using scRNA-Seq data of the corresponding normal tissue, removing TFs that are not specific of the epithelial cells of the tissue, for instance because they are specific to tissue-specific fibroblasts or endothelial cells. **d)** Subsequently, the regulons of each TF can be applied to a scRNA-Seq dataset comprising normal, precancerous and cancer-stages to infer for each cell and TF, the TF’s differentiation activity (TDA), resulting in a TDA-matrix defined over cells and TFs. This TDA-matrix can then be analyzed using diffusion maps to yield a 2D representation of a cellular ‘state-space’. **e)** To define the elevation in this landscape, representing cellular plasticity, we use a computationally fast approximation to diffusion entropy, called CCAT, which estimates the efficiency of intracellular signaling using a high-quality PPI network as scaffold. CCAT can be computed as the Pearson Correlation Coefficient (PCC) between a cell’s transcriptomic profile and the PPI’s degree profile **f)** Integrating the cellular plasticity with the 2D diffusion map representation of TDA-space allows construction of a Waddington landscape with attractors representing normal, precancer and cancer-states, and allowing identification of high cancer-risk states, as those proximal to the cancer-state. **g)** Validation of CCAT as a measure of cellular plasticity in early development, illustrating how CCAT decreases with developmental timepoint for 4 different cell-lineages. Total number of cells, and number of cells at each timepoint is indicated. P-values are from a linear regression test.

### Further validation of CCAT as a measure of cellular plasticity

Before applying PLASTIC to precancerous lesions, we decided to further demonstrate CCAT’s ability to measure plasticity in diverse and novel contexts, including mouse fetal development and drug resistance in longitudinally sampled data. First, we computed CCAT in 1.78 million cells encompassing 38 cell-types during embryonic mouse development (E9.5,10.5,11.5,12.5,13.5) [69] (**Methods**), which showed that CCAT decreased significantly with developmental timepoint in all 38 cell-types (**Fig.1g, SI fig.S4**). Second, if CCAT is a general measure of plasticity and not just a measure of developmental potential, it ought to also correlate with drug resistance. Analyzing a serial scRNA-Seq dataset profiling pretreatment, posttreatment and metastasis samples from the same breast cancer patient [70], confirmed that CCAT of epithelial cells increases with treatment and metastasis, and most importantly that it also correlated with malignant EMT-like metaprograms [71] within the pretreatment epithelial cell population (**SI fig.S5a-c**). This demonstrates that it marks the more plastic EMT-like cells that confer drug resistance [70]. Similar associations were also observed in serial pre and post-treatment samples of a chronic lymphocytic leukemia (CLL) patient [72] (**SI fig.S5d-e**): we observed how the B-cells of the CLL samples segregated by pre and post-treatment status, with a significant number of high CCAT cells displaying similarity to both groups (**SI fig.S5d**), suggesting that these are the cells that survive treatment. Moreover, ranking B-cells from the pretreatment phase according to their CCAT values displayed a strong positive correlation with how similar each pretreatment B-cell is to those found in the post-treatment sample (**SI fig.S5e).** Collectively, these data show that CCAT is a generic measure of cellular plasticity as it correlates with both developmental/differentiation potential and drug resistance. Consequently, with PLASTIC we have a formal computational framework for quantifying both epigenetic reprogramming (i.e. differentiation state) and cellular plasticity.

### Validation of PLASTIC in Barrett’s Esophagus

In order to validate PLASTIC we first considered the case of Barrett’s Esophagus (BE), a metaplasia where the squamous epithelial cells of the esophagus get replaced by columnar epithelial cells (enterocytes/goblets) normally found in the colon, and which generally precedes the development of esophageal adenocarcinoma [73, 74]. Thus, one would expect the esophageal specific TFs to display reduced differentiation activity in BE compared to normal esophagus, whilst colon-specific TFs should be increased in line with the emergence of colon-epithelial cells. To test this, we analyzed a scRNA-Seq dataset encompassing 2009 cells, derived from 4 BE patients, each with 4 samples taken from different anatomical locations that include normal esophagus, normal duodenum, normal stomach, as well as from the BE lesion itself (**SI fig.S6a**). Using Harmony [75, 76] to integrate the data from the 4 patients, revealed clusters that broadly speaking correlated with anatomical location (**SI fig.S6b**). Next, we estimated the TDA levels of the esophageal, colon and gastric specific TFs across the 2009 cells (**Methods**). Taking the average over the respective TFs, we confirmed that esophageal-TFs were most active in the normal esophagus, that gastric-TFs were most active in the normal stomach cells and that the colon-TFs displayed higher levels in BE compared to normal esophagus (**SI fig.S6c**), consistent with BE being a form of intestinal metaplasia. To confirm that these average patterns are seen at the level of individual TFs, we compared the TDA levels of esophageal and colon TFs between BE and normal esophagus (**SI fig.S6d**). Unique esophageal and colon-specific TFs clearly displayed a corresponding preference for reduced and increased TDA in BE, respectively, with TFs specific to both esophagus and colon displaying more ambiguous patterns (**SI fig.S6d-e**). Of note, these patterns were also recapitulated by considering differential expression, with differential expression and differential TDA exhibiting relatively strong agreement (R=0.73, **SI fig.S6f**). Overall, this validates PLASTIC and shows that it could be used to identify metaplasia or other differentiation skews in preneoplastic contexts.

### Reduced differentiation activity in gastric intestinal metaplasia correlates with plasticity

Gastric intestinal metaplasia precedes intestinal-type gastric cancer (IGC) and is characterized by the emergence of intestinal enterocytes and goblet cells [51]. To explore the relationship between epigenetic reprogramming, plasticity and cancer-risk in IGC development, we re-analyzed a scRNA-Seq dataset from 9 patients, representing the 4 main stages in IGC development: normal atrophic gastritis (NAG, 3 samples), chronic atrophic gastritis (CAG, 2 samples), intestinal metaplasia (IM, 4 samples) and early gastric cancer (EGC, n=1) (**Fig.2a**, **Methods**)[77]. All patients contributed one sample, except the EGC patient for which an IM sample was also taken. After QC, we obtained a total of 42,533 cells, with dimensional reduction and visualisation using Seurat [78] and tSNE [79] revealing clusters that correlated with cell-type and not patient (**Fig.2a**). Next, we estimated TDA-levels for the 47 previously validated stomach epithelial specific TFs in the normal epithelial and stromal cells of the NAG patients. The TDA of 42 of these 47 TFs (i.e. 89%) were significantly higher (nominal P<0.05) in the normal epithelial cells compared to stroma (**Fig.2b**), thus validating the robustness of these TF-regulons. Focusing next on the subset of 32,726 epithelial cells, we obtained clusters that broadly speaking correlated with the main gastric epithelial subtypes, including glandular mucus cells (GMCs), chief and pit mucus cells (PMCs), as well as the IM cell-types (goblets, enterocytes) and cancer cells (**Fig.2c**). Patterns of TDA between the normal gastric epithelial types were consistent: for instance, the pit mucous cell specific TF *FOXQ1* displayed higher TDA levels in the corresponding PMC cluster compared to GMCs (**Fig.2d**).

**Figure-2:**
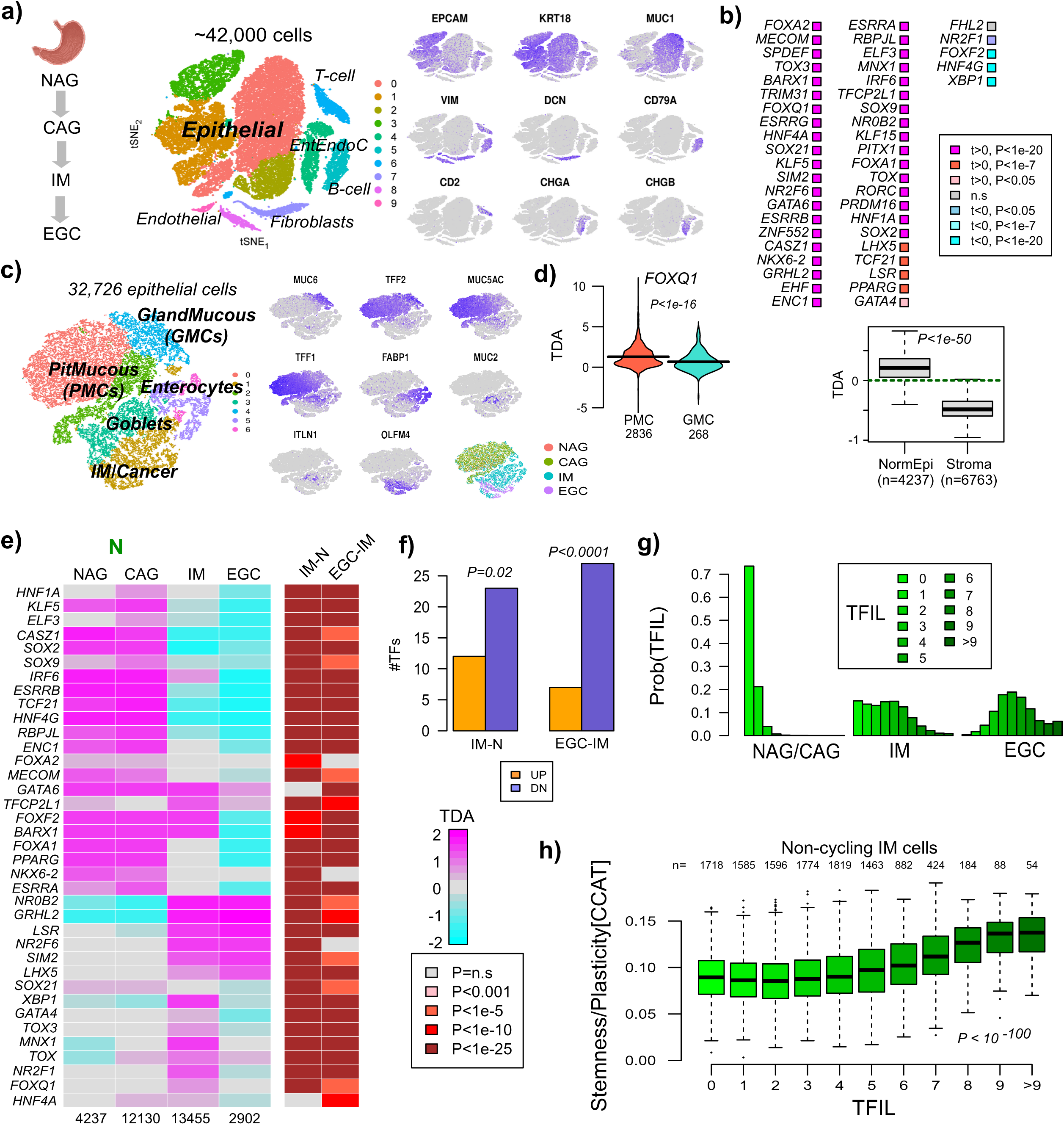
Dedifferentiation correlates with plasticity in IM preceding EGC. **a)** tSNE visualization plot of over 42,000 cells derived from 9 donors (3 with normal atrophic gastritis (NAG), 2 with chronic atrophic gastritis (CAG), 3 with intestinal metaplasia (IM) and 1 with matched IM and early gastric cancer (EGC)). Cells have been labeled according to cell-type, and expression levels of selected markers are shown in the subpanels on the right. **b)** Differential transcription factor differentiation activity (TDA) validation diagram for 52 stomach specific TFs, as assessed between the normal epithelial and stromal cells from the 3 NAG donors. P-values are from a two-tailed t-test. t labels the t-statistic. Boxplot below diagram compares the average TDA over all stomach specific TFs between the normal epithelial and stromal cells from NAG donors. P-value is from a one-tailed Wilcoxon rank sum test. Number of cells is given below boxplot. **c)** tSNE visualization plot for the epithelial cells only, with cells labeled by epithelial subtype, alongside panels labeling cells by the expression levels of known marker genes. Also shown is the distribution of cells according to disease stage. **d)** Violin plot comparing the TDA of *FOXQ1,* a pit mucus cell specific TF, between the pit mucus (PMC) and glandular mucus (GMC) cells derived from NAG donors. P-value is from a one-tailed Wilcoxon rank sum test. **e)** Heatmap of TDA levels of 37 stomach-epithelial specific TFs during progression from NAG+CAG to IM to EGC. The TDA values represent averages. Number of cells in each stage is indicated below heatmap. Color bars to the right indicate the statistical significance levels (two tailed P-values from a t-test) comparing IM to N (NAG+CAG), and EGC to IM. **f)** Barplot contrasting the number of significantly down (DN) and upregulated (UP) TFs between IM and N and between EGC and IM. P-values are from a one-tailed Binomial test to assess significance of skew. **g)** Barplots displaying the probability distribution of transcription factor inactivation load (TFIL) values for cells from the 3 different disease stages. **h)** Boxplot displaying the CCAT estimate of stemness/plasticity against the TFIL for all non-cycling IM-cells. P-value is from a linear regression.

Next, we explored the dynamics of TDA levels as epithelial cells progress from NAG and CAG to IM and EGC. Of 37 TFs that displayed higher TDA in normal gastric epithelial cells compared to gastric stroma (Bonferroni P<0.05), 23 displayed a reduction in TDA between IM and NAG/CAG, whilst 27 displayed a further decrease between IM and EGC (**Fig.2e**). Although the skew towards lower TDA in IM relative to NAG+CAG was only marginally significant (Binomial P=0.02, **Fig.2f**), this skew became more significant between the EGC and IM states (Binomial P<0.0001, **Fig.2f**), suggesting that the reduction in TDA is a trend that begins in the IM stage. Of note, this trend was not observed at the gene expression level, as the majority of stomach TFs displayed upregulation (**SI fig.S7a**), a pattern that is inconsistent with the reduced differentiation state of EGC, once again highlighting how mRNA TF levels are an unreliable indicator of a TF’s binding activity in differentiation. This is likely because the regulons of these TFs are rewired reflecting the acquisition of oncogenic functions, as observed and experimentally verified by us in other cancer-types like esophageal squamous cell carcinoma where pro-differentiation factors like TP63 and SOX2 metamorphose into oncogenes [45]. On the other hand, TFs displaying reduced expression with cancer progression generally did exhibit a reduction in TDA (**SI fig.S7b**). Importantly, unlike stomach differentiation TFs, colon-specific TFs did not display a preferential reduction in TDA, instead displaying a notable increase of TDA in the IM-stage (**SI fig.S8**), consistent with the emergence of metaplastic goblet and enterocyte cells (**SI fig.S9**).

Next, we reasoned that those cells in IM that display a higher number of inactivated stomach-TFs would be more stemlike and plastic and hence at higher risk of progression to EGC. To explore this, we computed the transcription factor inactivation load (TFIL) for each epithelial cell in the IM stage (**Methods**). This TFIL is obtained by counting the number of stomach-specific TFs for which the corresponding TDA levels are significantly lower in the cell compared to a null distribution determined by the average and standard deviation of the TDA levels over normal epithelial cells (NAG+CAG). In line with the previous analysis, IM displayed an increase in the TFIL, but importantly also a large spread of TFIL values, with EGC displaying a further increase but with a lower variance (**Fig.2g**). The larger variance of TFIL in IM compared to NAG/CAG and EGC, is consistent with the view that clonal heterogeneity is maximal just before cancer onset, with the emergence of cancer being characterised by selection of a more plastic stemlike state [32]. To test this, we estimated the plasticity level of each epithelial cell using CCAT. We observed a strong correlation between TFIL and CCAT among all IM cells, which remained strong when restricting to non-cycling IM cells (**Fig.2h**). Overall, these data indicate that IM cells with higher TFIL values (hence more dedifferentiated) also display increased plasticity.

### PLASTIC identifies a high cancer risk state characterized by loss of a fetal goblet gene expression program

Of the 13,455 epithelial cells derived from IM samples, there were 3333 enterocytes, 3626 goblets, 873 glandular mucus cells, 2277 pit mucous cells and 3346 cells that clustered closely to the cancer cells from the EGC samples (**Fig.2c**), which we henceforth term ‘precancerous’ (PreC). Using the estimated TDA levels, we next built a diffusion map [61] of all the cells from the IM stage, which revealed that the PreC cells seem to arise from either the goblet or enterocyte cells that define IM (**Fig.3a**), consistent with the Correa cascade model [52, 80]. Moreover, the TFIL dedifferentiation index was significantly higher for the PreC and IM-branch (goblet+enterocyte) cells (**Fig.3a**). Rederiving the diffusion map with the cancer cells from the EGC state included, further confirmed that the PreC cell state from the IM-stage is closest to the cancer-state (**Fig.3a**). Using the diffusion map embedding and the CCAT plasticity score, we next visualized a corresponding Waddington epigenetic manifold for the various cellular states (**Fig.3b**). Based on this manifold, we estimated the relative probability of each cell and cell-state to progress to EGC, thus defining a cancer progression risk score (CRS) (**Methods, Fig.3c**). This risk score was highest in the PreC and goblet cells (**Fig.3d**). Moreover, the risk score was strongly correlated with the TFIL, independently of cell-proliferation (**Fig.3e**). Put together, this suggests that the cancer clone defining EGC likely emerges from an aberrant goblet progenitor cell that has undergone further dedifferentiation.

**Figure-3:**
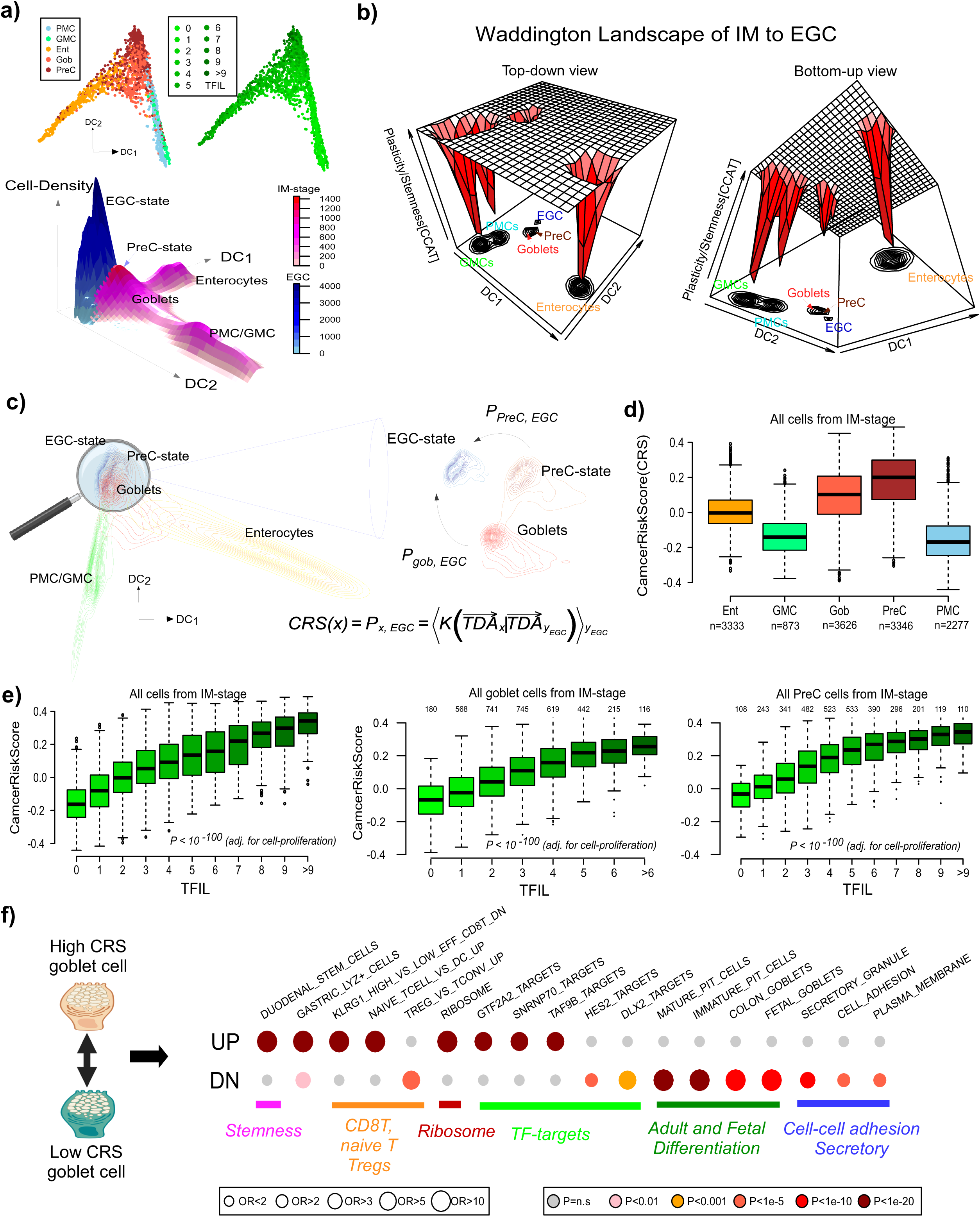
Waddington landscape of intestinal metaplasia to EGC progression. **a)** Diffusion map of all epithelial cells collected from IM samples, with cells labeled by cell-type and TFIL score. Lower 3D panel is a depiction of the diffusion map including now also the cells from the early gastric cancer (EGC) stage, with the elevation describing cell-density. To better highlight the EGC state, densities were computed separately for IM and EGC cells. **b)** Waddington landscape depiction of the main cell-states found in IM, including the EGC-state. The x and y dimensions are the diffusion components of panel a), whilst the z-dimension now measures the stemness/plasticity of the state as estimated by CCAT. The depth of the attractors is thus inversely proportional to plasticity/stemness. **c)** Graphical depiction of how the cancer-risk score of IM-stage cells is computed. Briefly, for each cell from IM-stage labeled *x* we compute a kernel *K* between the cell’s TDA profile and the TDA of each cancer cell from the EGC-stage, labeled *y.* Finally, an average is taken over all EGC cells *y,* to arrive at a relative cancer-risk-score (CRS) for each IM-stage cell *x, CRS(x).* **d)** Boxplot displaying the CRS across all cell-types from the IM-stage. Number of cells for each cell-type is given below. **e) Left:** Boxplot displaying the CRS of all IM-stage cells, with cells stratified by their TFIL (dedifferentiation score). Number of cells for each TFIL value is given below. P-value is from a linear regression of CRS vs TFIL adjusted for cell-proliferation. **Middle & right**: As left panel, but now restricting to goblet and precancerous (PreC) cells only. **f)** Gene Set Overrepresentation analysis of genes up and downregulated between high CRS/TFIL and low CRS/TFIL goblet cells from the IM-stage. Selected enriched terms from the Molecular Signatures Database are shown with their odds ratios (OR) and one-tailed Fisher test P-values of enrichment.

To explore this further we studied the cellular heterogeneity within the goblet and PreC cell populations that emerge within the IM-stage. Within both populations, the TFIL was also strongly associated with the CRS, independently of cell-proliferation (**Fig.3e**). Thus, even within the goblet cells that emerge during IM, there is heterogeneity that may indicate differential cancer risk. To gain support for this, we performed DE-analysis between high and low cancer risk IM goblet cells (**Methods**). There were approximately twice as many downregulated genes (n=620, FDR<0.001) than upregulated ones (n=313, FDR<0.001). Performing an overrepresentation analysis on the top 313 up and downregulated genes (**SI table S3**) revealed how upregulated genes were enriched for gastro-epithelial stemness and ribosomal genes. Consistent with this, ribosomal gene overexpression has been associated with stemness and multipotency across many different lineages and species and does so independently of cell-proliferation [32, 62, 67, 81–84]. In contrast, downregulated genes were notably enriched for extracellular terms including cell-cell adhesion and secretory granules, as well as goblet and gastric cell differentiation, including those characterizing fetal goblet cells (**Fig.3f**). A similar pattern of enrichment was also seen for genes differentially expressed between the high and low CRS/TFIL precancerous (PreC) cells (**SI fig.S10, SI table S4**). Using the DESCARTES scRNA-Seq database of human fetal development [85], we built our own fetal goblet gene expression signature (**Methods**, **SI table 5**) and verified that most of these genes were downregulated in the high cancer risk goblet and PreC states (**SI fig.S11**). The observation that genes expressed by fetal goblet cells and genes implicated in cell-adhesion are downregulated, suggests that the high CRS/TFIL goblet and PreC cells have stem or multipotent like features of early developmental cells that precede the fetal stage. Overall, these results indicate how PLASTIC is able to identify a highly dedifferentiated and plastic goblet progenitor state that closely resembles the cells found in EGC.

### PLASTIC identifies a high cancer risk state in colonic polyps

Next, we considered a different form of cancer precursor lesion, namely colonic polyps, within the context of colon adenocarcinoma progression [53]. We analyzed a snRNA-Seq dataset encompassing over 200,000 cells derived from 8 donors with Familial Adenomatous Polyps (FAP) and 7 non-FAP ones, comprising 81 samples including 27 normal tissues (N), 48 polyps (P) and 6 colorectal adenocarcinomas (CRC) [53]. Focusing first on the normal samples (39,590 normal cells) (**Fig.4a**), we verified that all of our 18 colon-epithelial TFs displayed higher differentiation activity in the normal enterocytes and goblets compared to immune, endothelial and fibroblast cells (**Fig.4b**). Had we used mRNA expression, 16 of these would be upregulated in enterocytes and goblets (**Fig.4b**). Next, we focused on the enterocytes (n=104,009) from all stages in colorectal adenocarcinoma progression, estimating TDA levels of the 18 colon-epithelial specific TFs in each cell. Average TDA over these 18 TFs displayed a clear gradual decrease, from normal to polyps to colorectal adenocarcinoma (**Fig.4c**). Of these 18 TFs, 15 displayed reduced differentiation activity in polyps compared to normal colon epithelial cells (**Fig.4c**), a highly significant skew (one tailed Binomial test P=0.0006). For each of the 65,285 cells from polyps (and using the normal epithelial cells as reference), we computed their TFIL. We also computed the similarity of each polyp cell’s TDA profile to those of cancer cells from the CRC stage, thus defining a separate cancer-risk score (CRS). Remarkably, the TFIL and CRSs were strongly correlated, an association that was also independent of cell-proliferation (**Fig.4d**). Thus, once again, the preneoplastic cells most similar to cancer cells (therefore assumed to be at higher risk for selection), can be identified as those having a higher number of inactivated tissue-specific TFs.

**Figure-4:**
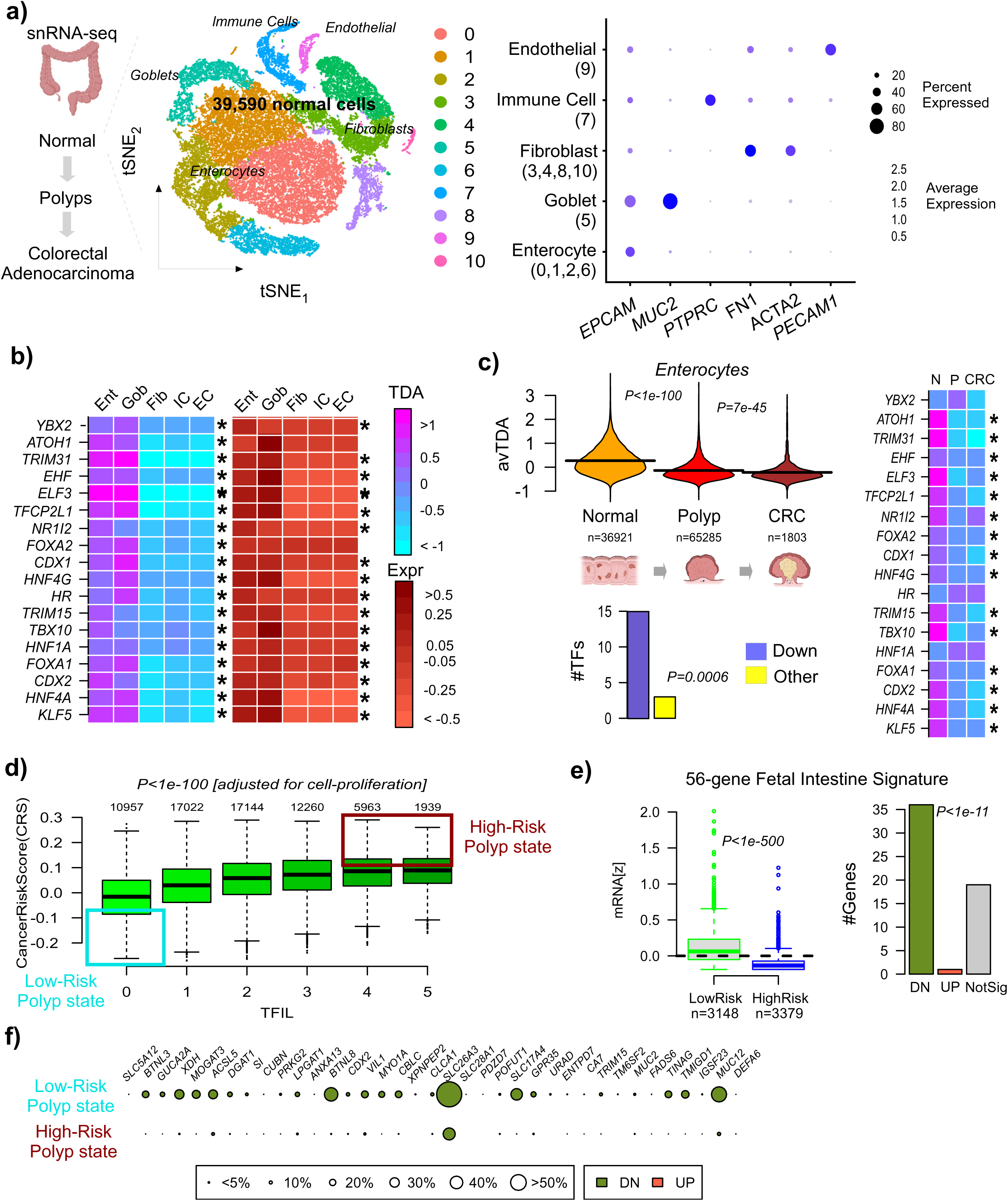
PLASTIC identifies a high cancer risk polyp state characterized by loss a fetal intestine gene expression program. **a)** tSNE plot depicting 39,590 cells from healthy donors, with cells annotated by Seurat cluster and cell-type. Bubble plot to the right justifies the labeling of clusters (integer numbers below cell-type names) by cell-type, according to their expression patterns over selected well-known marker genes. **b)** Heatmap of estimated TDA levels (left) and mRNA expression levels (right) of 18 colon-epithelial-specific TFs. * Means significantly higher TDA levels in enterocytes (Ent) and goblets (Gob) compared to each of immune (IC), fibroblast (Fib) and endothelial (EC) cells. Bonferroni P<0.05. **c)** Violin plot of the average TDA level over 18 validated colon-epithelial specific TFs with cells stratified by disease stage, with the number of cells in each stage indicated below the plot. P-values are from Wilcoxon rank sum tests comparing Normal to Polyp and Polyp to CRC stages. Heatmap to the right depicts the average TDA values for the 18 validated colon-epithelial-specific TFs across the 3 states. *Bonferroni P<0.05. Barplot beneath violin plot compares the number of such TFs that display reduced TDA during progression to polyp and CRC stages to TFs not displaying such a reduction. P-value is from a one-tailed Binomial test. **d)** Boxplot of the cancer risk score (CRS) against TFIL for all polyp cells. Number of cells in each TFIL category is given. P-value is from a linear regression adjusted for cell-proliferation. Highlighted boxes indicate our definition of high and low cancer-risk polyp states. **e)** Boxplot comparing the average expression of a 56 gene fetal intestine signature from DESCARTES, between the high and low risk polyp states. P-value is from a one-tailed Wilcoxon test. Barplot to the right quantifies the number of genes from this signature that are significantly downregulated/upregulated (FDR<0.05) or not significantly altered. P-value is from a one-tailed Binomial test assessing skew towards downregulation. **f)** For the differentially expressed genes in e), balloon plot displays the percentage of cells expressed by each gene in each of the low and high risk polyp states. Size of balloons indicates percentage of expressed cells.

### High cancer risk state in polyps displays loss of a fetal intestine gene expression program

To explore the biological significance of this high-risk polyp state, we performed a differential expression analysis between polyp cells with a high TFIL and CRS against those with a low TFIL and CRS. Of the 8610 differentially expressed genes (Wilcoxon test adjusted P-value < 0.001), the overwhelming majority were downregulated in the high-TFIL/risk group, with only 398 genes displaying upregulation. Performing Gene Set Overrepresentation Analysis (**Methods**) [86, 87] on the top 398 upregulated and top 398 downregulated genes (**SI table S6**), revealed top-ranked enriched terms for upregulated genes that mapped to stem-cells as characterized in the duodenum, as well as gastric isthmus cells and large intestine MKI67+ cells (**SI fig.S12**). We also observed enrichment for ribosome, translation and spliceosome GO-terms (**SI fig.S12**), similar to what we found in the high cancer-risk IM state, and consistent with previous studies [62, 66, 88]. Among both up and downregulated genes, there was enrichment for genes associated with T-regulatory function and CD8+ T-cell exhaustion. Among downregulated genes, enriched terms mapped to cell-adhesion and cadherin binding, and to genes normally expressed in mature enterocytes, pit cells and fetal intestinal epithelial cells (**SI fig.S12**). Using the DESCARTES database to define a 56 gene signature specifically overexpressed in fetal intestinal epithelial cells (**Methods, SI table S5**), we verified that their average expression is significantly lower in the high-risk polyp-state, with 36 of 37 differentially expressed genes being downregulated (**Fig.4e-f**). This last finding indicates that our high-risk polyp state defines a highly dedifferentiated state, even when compared to fetal intestinal cells. Thus, once again, from a normal developmental perspective, the high cancer-risk state is less differentiated than the corresponding fetal epithelial state.

### PLASTIC predicts dedifferentiation and squamous metaplasia in lung cells from smokers

Next, we applied PLASTIC to a 10X scRNA-Seq dataset profiling lung tracheal epithelial cells from 6 healthy heavy smokers and 6 age-matched healthy never-smokers [89] (**Methods, Fig.5a**). We reasoned that PLASTIC would enable us to determine if smoking is associated with a conversion of lung basal cells to squamous metaplasia, as squamous metaplasia normally precedes lung squamous cell carcinoma development, the lung cancer-type for which smoking is also the major risk factor [90, 91]. First, we aimed to identify lung epithelial TFs that are specific to each lung epithelial cell-type, including the 3 main subtypes, i.e. basal, cilia and goblets. This was done by first processing the 14,435 cells from healthy non-smokers only, which revealed basal (n=7731), cilia (n=1103) and goblet (n=1000) clusters (**Fig.5a**). We next estimated the TDA for 38 lung-specific TFs across all these epithelial cells (**Methods**). PCA over the TDA-matrix revealed how PC1 discriminates cilia from goblets and basal cells, whilst PC2 was able to discriminate the different basal subtypes from each other, notably differentiated vs proliferating (**Fig.5b**). Of the 38 lung-specific TFs, only one was specific to cilia (*FOXJ1*), 10 to goblets (*TAL1, TBX4, GATA2, LDB2, ELF3, SOX18, XBP1, LSR, PPARG* and *NR2F2*), 19 to basals (*TBX2, EPAS1, ETV1, ERG, AHR, PML, BGN, ZFP36, TNXB, TEAD2, MEOX2, KLF4, HIF3A, KLF9, STON1, ZFP36L2, CEBPD, TRIP10* and *TGFB1I1*), 4 specific to both cilia and goblets (*FOXA1, FOXA2, NKX2-1* and *TFEC*) and 4 specific to goblets and basals (*EHF, AFDN, SOX13, ETS1*) (**Fig.5c**). Next, we re-analyzed the data including now the 15,578 cells from heavy smokers, which led to a similar clustering as before, with clusters driven by cell-type and not by donor ID or smoking status (**SI fig.S13**). To explore if smoking affects the lung cell-type specific TFs, we re-calculated the TDA over all 30,013 cells, comparing TDA levels of smokers vs non-smokers, stratified by the 3 main cell-types. Of the five cilia-activated TFs, four displayed a significantly lower TDA in the cilia of smokers vs non-smokers, including the cilia-specific factor *FOXJ1* and the cilia-goblet specific ones (*FOXA1, FOXA2, NKX2-1*) (**Fig.5d**). The reduced TDA of *FOXJ1* in the cilia of smokers is consistent with reports of shorter cilia in smokers [92]. Of note, ordinary differential expression analysis would have failed to predict this (**SI fig.S14**), highlighting once again how TDA is more reliable than DE. Of the 18 goblet-activated TFs, a total of 8 displayed reduced TDA in goblets from smokers, which included once again *NKX2-1* as well as *GATA2, SOX18* and *TBX4;* in basal cells, 12 of the 23 basal-TFs displayed reduced TDA in smokers (**Fig.5d**). Overall, as assessed across all 3 cell-types, a total of 24 TFs displayed reduced TDA in smokers, whilst only 9 displayed increased TDA (**Fig.5d**), representing a significant skew towards reduced differentiation activity (Binomial one-tailed P=0.002). This indicates that smoking is preferentially associated with a reduced differentiation activity of cell-type specific TFs in each of the 3 lung cell-types. To assess if smoking is associated with a transformation to squamous metaplasia, we considered our earlier TFs that are specific to the esophagus, since the esophagus epithelium is of the squamous subtype. Thus, we calculated the TDA of all 43 esophageal-specific TFs, which revealed that in each of the 3 lung epithelial cell-types there was a clear skew towards increased TDA in smokers, consistent with an increase in squamous differentiation (**Fig.5e,** Binomial one-tailed P=0.0002). We verified that key squamous differentiation TFs like *PAX9, TP63* and *BARX2* displayed increased TDA in basal cells from smokers (**Fig.5e**), consistent with the emergence of squamous metaplasia.

**Figure-5:**
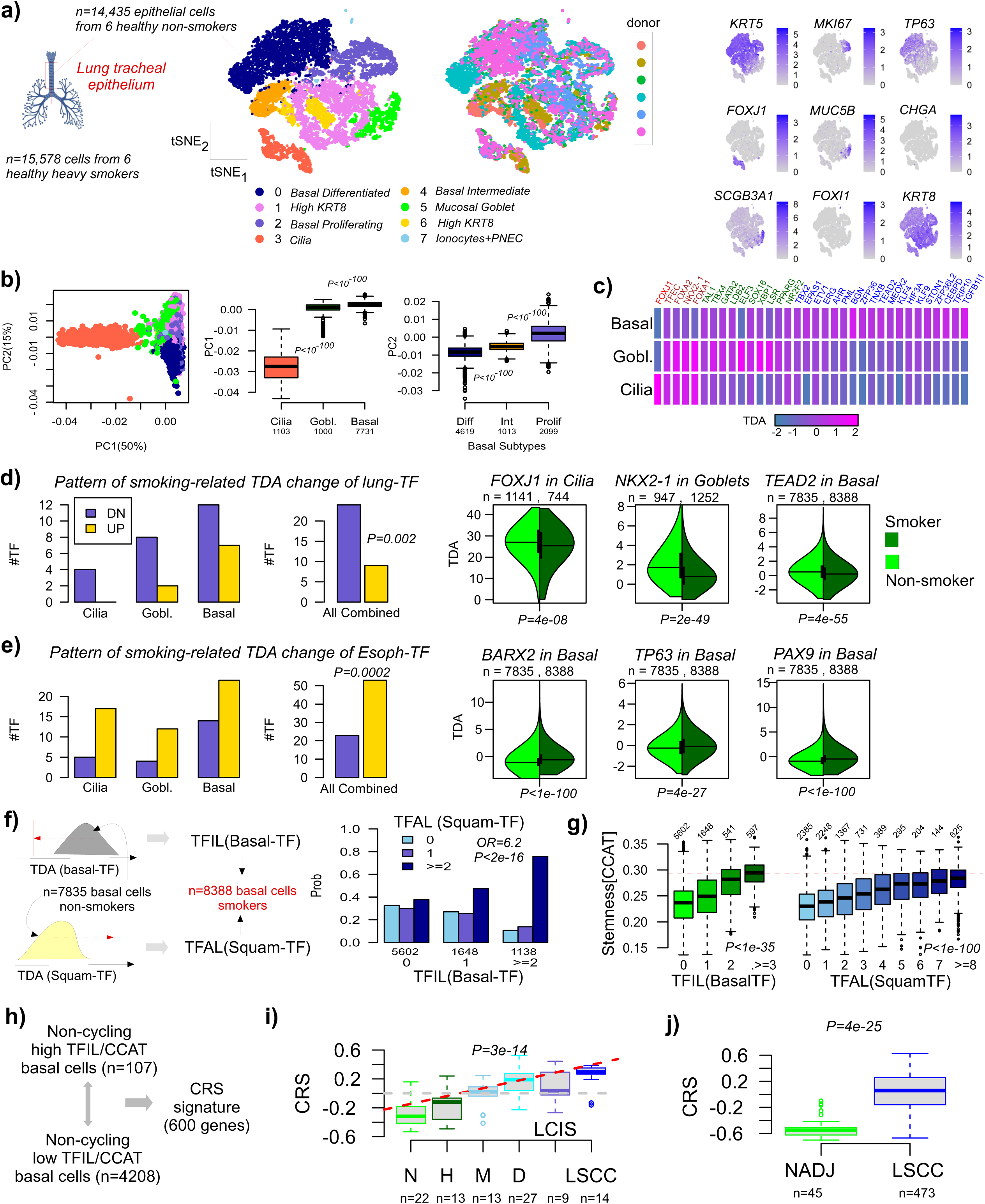
PLASTIC reveals squamous metaplasia and a high cancer risk state in basal cells from smokers. **a)** tSNE plots of 14,435 lung tracheal epithelial cells from healthy non-smokers with cells annotated by cluster (left), donor ID (middle) and marker expression (right panels), justifying the labeling of clusters by cell-type. **b) Left:** PCA scatterplot where PCA was performed over a transcription factor differentiation activity (TDA) matrix defined over the 14,435 lung epithelial cells from non-smokers and 38 lung-epithelial specific TFs. **Middle:** Boxplot of PC1 vs cell-type, stratified by basal, goblet and cilia subtypes. P-value is from a two-tailed Wilcoxon rank sum test comparing the two cell-types on either side of the P-value. **Right:** Boxplot of PC2 vs cell-type, focusing on only the 3 basal subtypes. P-value is from a two-tailed Wilcoxon rank sum test comparing the two cell-types on either side of the P-value. **c)** TDA heatmap of lung epithelial specific TFs displaying their average TDA in basal, cilia and goblet cells from non-smokers. TFs have been colored according to whether they are only cilia specific (red), cilia and goblet specific (brown), only goblet specific (green) and only basal specific (blue). **d)** Barplot displaying the number of significantly activated and inactivated cilia, goblet and basal specific TFs in the corresponding cilia, goblet and basal cell-types from smokers compared to non-smokers. Right barplot is for all cell-types combined. P-value is from a one-tailed Binomial test. Violin plots display examples of the TDA profiles for 3 TFs in corresponding cell-types, comparing the TDA-values in smokers vs non-smokers. Number of cells of the given cell-type from smokers and non-smokers are given above each violin plot. Below the plot, we give the two-tailed P-value of a t-test. **e)** As d), but for the TDA of 43 esophageal (squamous) specific TFs. **f)** For each of the basal cells from smokers, we compute a transcription factor inactivation load (TFIL) which counts the number of basal-specific TFs that are inactivated in smokers vs non-smokers. Similarly, for the same cells we compute a transcription factor activation load (TFAL) which counts the number of squamous-specific TFs that are activated in smokers vs non-smokers. Barplot compares the distribution of basal cells from smokers according to their TFAL load, when stratified by 3 different levels of TFIL load. Odds Ratio (OR) and one-tailed Fisher-test P-value comparing TFIL=0 to TFIL>=2 groups are given. **g)** Boxplots display the stemness CCAT index vs TFIL(basal-TFs) (left) and TFAL (squamous-TFs) (right) for all basal cells from smokers. P-values are from a one-tailed linear model adjusted for cell-proliferation. **h)** Derivation of a cancer-risk signature by comparing expression of high TFIL to low TFIL basal cells. **i)** Boxplots display the cancer-risk-score (CRS) as evaluated in an Agilent gene expression dataset encompassing 98 lung tissue samples encompassing normal (N), hyperplasia (H), metaplasia (M), dysplasia (D), lung carcinoma in-situ (LCIS) and invasive LSCC. P-value is two tailed from a linear regression. **j)** As i), but in the TCGA LSCC dataset. P-value is from a one-tailed Wilcoxon test.

### PLASTIC identifies a highly dedifferentiated plastic cancer-risk state in smokers

Focusing next on basal cells from smokers, we asked if cells with a higher number of inactivated basal specific TFs also display a corresponding higher number of activated squamous epithelial-specific TFs (**Fig.5f**). We found that basal cells from smokers with no inactivated lung basal TFs displayed a roughly equal probability of having 0, 1 or more than 2 activated squamous-TFs, whilst basal cells with at least 2 inactivated basal TFs displayed a significantly higher probability of also having 2 or more activated squamous TFs (**Fig.5f**, OR=6.2, Fisher one-tailed P<2e-16). This suggests that conversion to squamous metaplasia is intricately linked to a reduced basal differentiation activity. To explore if basal dedifferentiation and squamous differentiation (as measured by the number of inactivated basal-TFs and activated squamous-TFs, respectively) relates to stemness and plasticity, we computed the CCAT-index [32, 62, 66] across all basal cells from smokers, which revealed strong associations with plasticity, which were independent of cell-proliferation (**Fig.5g, Methods**). Of note, the group of basal cells from smokers displaying the highest plasticity were marked by cells with more inactivated basal TFs, as opposed to more activated squamous TFs (**Fig.5g, SI fig.S15a**), suggesting that the acquisition of stemness/plasticity is determined mainly by the reduced activity of basal-TFs. To shed further insight into the biological characteristics of this higher plastic/stem-like state, we performed differential expression and overrepresentation enrichment analysis comparing basal cells from smokers with high TFIL to corresponding low TFIL cells, restricting and not restricting to non-cycling cells (**Methods**). When restricting to non-cycling cells, we observed strong enrichment for oxidative phosphorylation, aerobic mitochondrial respiration and ATP production among upregulated genes, whilst hypoxia, TNF alpha signaling and apoptosis (p53 inactivation) were downregulated (**SI fig.S15b**). Lung basal differentiation was strongly enriched among downregulated genes, consistent with the increased TFIL values and reduced differentiation (**SI fig.S15b**). Of particular interest was also the observation of enriched pan-cancer metaprograms representing stressed cells [71], among downregulated genes (**SI fig.S15b**). Altogether, these results suggest that high TFIL basal cells from smokers display increased oxidative phosphorylation, ATP production and plasticity, whilst evading apoptosis via p53 inactivation, thus creating an environment conducive for accumulating genetic and epigenetic alterations. To demonstrate that this state is potentially informative of cancer-risk, we built a corresponding expression signature (**Methods**, **SI tables S7-S8**), which we then applied to a series of lung tissue samples (n=98) encompassing normal (n=22), hyperplasia (n=13), metaplasia (n=13), dysplasia (n=27), lung carcinoma in-situ (n=9) and invasive LSCC (n=14) [93], to estimate for each sample a cancer-risk-score (CRS) (**Methods**). Remarkably, this CRS increased during progression to LSCC, with specially pronounced increases between the hyperplasia and metaplasia stages, between metaplasia and dysplasia, and finally between dysplasia/LCIS and LSCC (**Fig.5h**). When applied to the TCGA-LSCC dataset [94], the CRS-score was also elevated in LSCC compared to normal-adjacent tissue (**Fig.5i**). Overall, these results highlight how PLASTIC is able to identify a high cancer risk state by dissecting the heterogeneity of lung basal cells from smokers.

### Loss of a fetal basal expression program underpins progression to LSCC

Unlike intestinal metaplasia of the stomach or adenomatous polyps, we did not observe a downregulated fetal lung expression program in the high cancer risk state of smokers. This could be due to the early nature of squamous metaplasia in these samples, as these specimens were collected from heavy smokers that were otherwise healthy, i.e. without an explicit diagnosis of squamous metaplasia. Hence, to explore this further, we assembled 4 fetal lung expression signatures from the DESCARTES scRNA-Seq database [85] (**Methods, SI table S5**). These signatures represent genes that are specifically overexpressed in squamous, cilia, alveolar and basal cells from fetal lung tissue. Applying them to the earlier series of 98 lung-tissue samples confirmed increased expression of the squamous signature in metaplasia compared to normal and hyperplasia samples (**Fig.6a**), a pattern also evident by inspecting mRNA or TDA levels of squamous epithelial TFs (**Fig.6b**). The fetal squamous signature was downregulated in LSCC compared to LCIS, dysplasia and metaplasia stages, but still remained higher compared to normal and hyperplasia samples (**Fig.6a**), consistent with the cancer samples being diagnosed as LSCC. In contrast, the fetal basal signature displayed a significant downregulation in the LSCC state compared to all previous stages (**Fig.6a**). Hence, this suggests that loss of a fetal basal gene expression program also underpins progression to LSCC, consistent with our earlier finding that increased plasticity in basal cells is mainly driven by high basal TFIL scores and not high squamous TFAL ones. Further supporting this, the mRNA and TDA levels of basal-specific TFs displayed a progressive decrease during LSCC progression (**Fig.6c**). It is noteworthy that a much higher proportion of basal-specific TFs displayed a decrease in TDA during LSCC progression, compared to a relatively lower proportion of fetal lung basal-cell expressing genes displaying downregulation (**Fig.6d-e**), pointing once again to the importance of focusing on the dynamics of TFs. Overall, these data highlight how basal-specific TFs become inactivated during LSCC progression, with a corresponding loss of a fetal basal gene expression program between the non-invasive and invasive stages.

**Figure-6:**
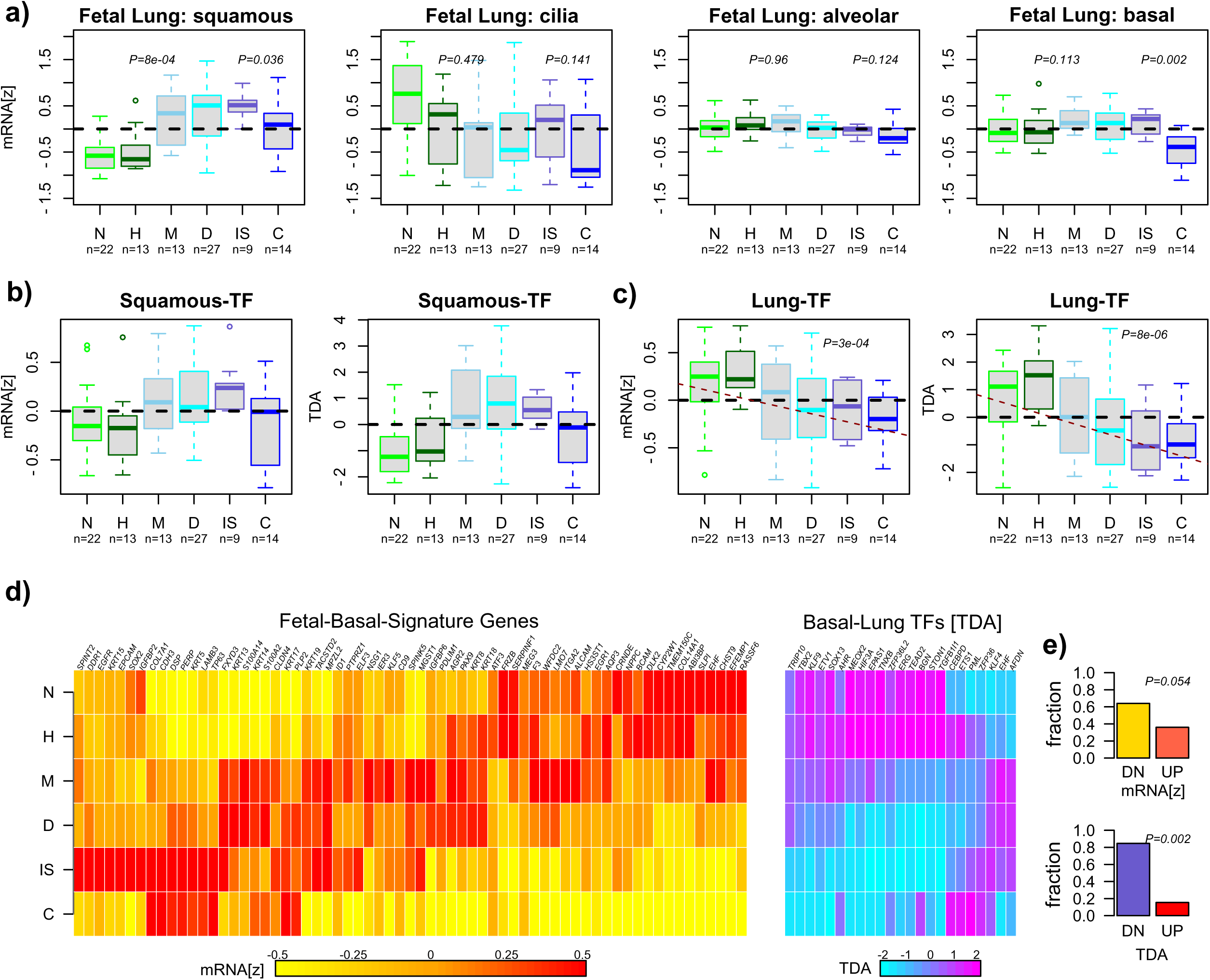
Dynamics of fetal gene expression during LSCC progression. **a)** Boxplots display the average mRNA expression (z-score transformed expression values) of 4 fetal lung gene expression signatures in an mRNA expression dataset of multi-stage LSCC progression (N=normal, H=hyperplasia, M=metaplasia, D=dysplasia, IS=lung carcinoma in-situ, C=lung squamous cell carcinoma). Number of samples is indicated below each box. P-values are from one-tailed Wilcoxon rank sum tests comparing adjacent stages, as shown. **b)** As a) but displaying the average mRNA expression and average TDA levels of 43 squamous specific TFs. **c)** As b) but for 23 lung basal specific TFs. Linear regression P-values and regression line are shown. **d)** Heatmaps of relative expression change for the fetal basal lung signature (right) and of the relative TDA change for lung-basal specific TFs. **e)** Barplots compare the relative fraction of genes from the fetal lung basal signature (top) and of lung basal TFs (bottom) that are significantly (Bonferroni P < 0.05) down and upregulated during LSCC progression. P-value is from a one-tailed Binomial test to assess significance of skew towards downregulation.

## Discussion

Epigenetic reprogramming and increased cellular plasticity have emerged as two enabling cancer hallmarks that are thought to play a particularly important role in the stages that precede cancer development. Although scRNA-Seq data of precancerous lesions offers an unprecedented opportunity to explore the role of epigenetic reprogramming and cellular plasticity in cancer development, a major challenge is how to quantify these features in such data. Our work is a major advance in presenting and extensively validating a pan-tissue computational framework (PLASTIC) that allows direct quantification of epigenetic reprogramming and cellular plasticity in normal, precancerous and cancer samples.

Our metric to quantify epigenetic reprogramming is anchored on estimating the differentiation activity of relevant lineage-specific TFs, which, as shown here and in our previous studies [44, 50], significantly outperforms differential expression (DE) as a means of assessing differentiation activity. This is a key point worth emphasizing further: the mRNA expression level of a TF in scRNA-Seq (and bulk RNA-Seq) data is often not a good proxy for TF-binding activity in relation to the TF’s function in differentiation. Our regulon-based approach, which is anchored on measuring the mRNA expression levels of direct and indirect downstream targets that mediate a TF’s function in differentiation, is a much better proxy of the TF’s activity in effecting differentiation, thus enabling a much more accurate estimation of the differentiation state of cells. Moreover, we have previously shown how lineage-specific TFs may metamorphose into oncogenes during oncogenesis (e.g. SOX2 and TP63 in ESCC), with these TFs displaying increased mRNA expression, yet displaying an altered cistrome, i.e. these TFs no longer bind at genes that effect differentiation (our SCIRA-regulons) but at other genes that map to cell-cycle and cell-proliferation pathways [45]. Thus, focusing on the mRNA levels of lineage-specific TFs could be misleading. Instead, by estimating the TDA of lineage-specific TFs we can more accurately chart the differentiation state of cells and quantify the level of epigenetic reprogramming they have undergone, irrespective of whether this reprogramming is caused by somatic mutations, copy-number changes or epigenetic alterations.

Importantly, we have seen how we can combine our TDA-estimates of lineage-specific TFs with a completely independent measure of cellular plasticity to construct Waddington landscape proxies of the precancer-state. To our knowledge, this is one of the first attempts to build such attractor landscapes from real scRNA-Seq data. As illustrated in the context of intestinal metaplasia that precedes IGC, this landscape is predictive, as it allows direct correlation of inferred precancer-states in the metaplasia to the invasive cancer one, thus pointing towards the more likely cell-of-origin for IGC. Whilst our finding that IGC may arise from an aberrant dedifferentiated goblet cell may appear surprising, it is in our opinion entirely consistent with the well-known Correa model [52]. Indeed, it is worth noting that we were able to characterize extensive heterogeneity within the IM goblet cell population, identifying a subset with increased stem-like features and downregulation of genes normally expressed in fetal goblet cells. This points towards the cell-of-origin of IGC being an aberrant goblet progenitor cell that in a normal physiological and developmental setting would precede and gives rise to goblet cells in the fetal stomach. Indeed, whilst goblet cells are absent from adult normal stomach, they are an important cell-type in the context of fetal stomach development [85], which then re-emerges in the context of IGC development.

Using PLASTIC we have also clearly demonstrated how smoking leads to a loss of dedifferentiation in each of the 3 main tracheal epithelial cell-types (cilia, basal and goblets), whilst in basal cells it also predicts increased activity of TFs implicated in squamous differentiation, consistent with the initial stages of squamous metaplasia. Importantly, within the basal cell population from smokers, PLASTIC identified a high cancer risk state, which in a multi-stage dataset representing LSCC development, displayed progressive and pronounced increases, not only between hyperplasia and metaplasia, but also between metaplasia and dysplasia, and between dysplasia/LCIS and invasive LSCC. Moreover, we found that within the basal cell population, a higher basal-TFIL led to a more pronounced increase in plasticity compared to a higher squamous-TFAL, suggesting that basal cells that have undergone more pronounced metaplastic transformation into a squamous subtype have a lower cancer-risk than those with more inactivated basal-TFs.

More generally, PLASTIC has revealed a common feature shared between the progression paths to all 3 cancer-types (IGC, LSCC and COAD): in the case of IGC and COAD, the high cancer risk state in IM and polyps, respectively, was characterized by a high level of dedifferentiation and a loss of a fetal gene expression program (a fetal goblet program in IM and a fetal enterocyte program in polyps); in the case of progression of squamous metaplasia and dysplasia to LSCC, it is loss of a fetal lung basal expression and basal-TF program that associates with progression to invasive LSCC. Thus, in all cases, PLASTIC points towards the high cancer risk state presenting an aberrant cellular state that, from a normal developmental perspective, appears to precede the fetal stage. This is consistent with the increased plasticity displayed by these high cancer risk states, and is an observation of profound significance, as other recent studies, using more traditional computational methods, have not elucidated this putative principle of oncogenesis [95].

Finally, it is important to stress that the lineage-specific TFs that orchestrate fetal and adult tissue development, and which we have used here to quantify the level of epigenetic reprogramming, are distinct from classical tumor suppressors like *TP53* or *RB1*. Whilst many of our lineage-specific TFs include known tumor suppressors (e.g. *ATOH1, CDX1, CDX2* in colon cancer [96–98], *NKX2-1*, *TBX4* in lung cancer [99, 100]), *GATA4, PITX1* in gastric cancer [101, 102]), a key distinction from classical tumor suppressors is in relation to their function and how they are preferentially inactivated. As shown by the TCGA project, classical tumor suppressors undergo frequent inactivation via somatic mutations or genomic deletions, with their inactivation targeting mostly apoptosis, cell-cycle and cell proliferation pathways. Thus, their inactivation directly translates to increased cellular proliferation. In contrast, lineage-specific TFs are generally speaking not hotspots of genetic dysregulation, instead displaying a preference for aberrant promoter hypermethylation [103]. Indeed, as shown by us in the context of ESCC development [45], and also more generally [16, 104], lineage-specific TFs undergo preferential promoter hypermethylation in normal cells as a function of exposure to diverse cancer risk factors, including age [105], smoking [106], estrogen [18] and viral infections [17]. Thus, it is very likely that the source of epigenetic reprogramming that we detect in the precancerous lesions of solid tissues is mostly of epigenetic origin. Moreover, our data supports the view that such epigenetic reprogramming is intricately linked to an increased cellular plasticity.

In summary, we have presented and extensively validated a computational framework for quantifying epigenetic reprogramming and cellular plasticity of single cells from scRNA-Seq or snRNA-Seq data, and in the context of different precancer lesions, including dysplasia and various forms of metaplasia. In each case, PLASTIC identified an epigenetically reprogrammed high cancer risk state characterized by increased plasticity and loss of a fetal gene expression program. Once scRNA-Seq/snRNA-Seq is cost-effective for clinical application, PLASTIC’s pan-tissue framework will have significant translational potential for developing cancer risk assessment and cancer preventive strategies.

## Methods

### Single-cell and single-nucleus RNA-Seq datasets

For validation and testing purposes of PLASTIC, we considered the following scRNA-Seq/snRNA-Seq datasets:

*1) Barrett’s Esophagus (BE):* The single-cell count Smart-Seq2 data was obtained from the Supplementary Information document as provided by the original publication [73]. For our analysis, we selected cells from four distinct anatomical locations (Normal Esophagus, Gastric Cardium, Duodenum and Barrett’s Esophagus) from each of 4 BE patients (PtA, PtB, PtC, PtD). Because the provided count data contained some NA-values, we only selected genes with no NA’s across all cells. After processing and QC with Seurat [78], we were left with a normalized scaled data matrix defined over 26733 Ensembl genes and 2009 cells. Because number of cells was evenly distributed across anatomical locations and patients, data was batch-corrected using Harmony [76]. We subsequently used the Harmony output to also build a harmonized data matrix over 22,024 protein coding genes (NCBI identifiers) and 2009 cells (450 normal esophagus, 622 gastric, 349 duodenum and 588 BE). This matrix was used as input for all follow-up analyses. To test robustness, all analyses were also performed on Seurat-normalized non-Harmony adjusted data, yielding very similar results.
*2) Intestinal type gastric cancer (IGC) progression:* We downloaded the single cell count matrices from GEO (GSE134520) [77]. This 10X scRNA-Seq dataset is derived from 9 patients, representing the 4 main stages in the development of IGC: normal atrophic gastritis (NAG, 3 samples), chronic atrophic gastritis (CAG, 2 samples), intestinal metaplasia (IM, 4 samples) and early gastric cancer (EGC, n=1)[77]. All patients contributed 1 sample, except the EGC patient for which an IM sample was also taken. After processing and QC with Seurat, we obtained a normalized data matrix defined for 18107 genes and 42,533 cells. Of these, a total of 32,724 are epithelial. The final matrix defined over NCBI gene IDs and epithelial cells was of dimension 14,334 genes times 32,724 epithelial cells: 4237 cells from NAG, 12130 cells from CAG, 13455 from IM and 2902 from EGC.
*3) Colon adenocarcinoma progression:* We analyzed a 10X snRNA-Seq dataset of colon adenocarcinoma progression, including normal, polyp and adenocarcinoma states from Becker et al [53], which included people with and without familial adenomatous polyps (FAP). We used the normalized form of this dataset, as processed and described by us previously [107]. Briefly, we initially used Seurat to process the normal cells (n=39,667) from healthy non-FAP individuals, in order to identify cell-types and to build a cell-type annotation predictor using the built-in procedure within EpiSCORE [108]. This predictor was constructed at the resolution of 5 cell-types (enterocytes, goblets, immune cells, stromal and endothelial), because these formed well-defined clusters. We then processed the full dataset of 826,250 cells with Seurat resulting in 380,527 high quality cells, which were then annotated using our predictor. We only assigned cells to a given cell-type if the assignment probability was >0.7, resulting in a final normalized data matrix defined over 27,133 genes and 380,527 cells, including confidently assigned endothelial cells (n=1866), enterocytes (n=104,009), goblets (n=78,451), immune cells (n=24,973) and stromal cells (n=7941). The distribution of enterocyte cells between stages was 36,921 normal cells (17,795 from non-FAP + 19,126 from FAP-cases), 65,285 polyp cells (65,049 from FAP-cases) and 1803 cells from colorectal cancer (1663 from non-FAP cases).
*4) Lung squamous metaplasia and smoking:* We downloaded the processed Seurat R-object file from GEO (GSE134174) [89]. This 10X study sequenced a total of 68,370 epithelial cells (36,248 after QC and doublet removal) from 15 human tracheal healthy donors (6 heavy smokers, 6 never-smokers, 2 light smokers, and 1 pediatric subject). The data from the pediatric subject as well as that of the two light smokers. who were also younger than the never and heavy smokers, were excluded when drawing inferences about the effect of smoking, so as to avoid confounding by age.

We also analyzed two longitudinal scRNA-Seq datasets representing serial sampling of pre-and-posttreatment samples, of one breast cancer and on chronic lymphocytic leukemia (CLL) patient, in order to demonstrate that CCAT can identify drug resistant cells in the pretreatment population:

*5) Serial sampling of breast cancer patient:* This Chromium 10X scRNA-Seq longitudinal dataset is available from GEO GSE264205 [70]. We downloaded the raw count files from the GEO site and processed it using the Seurat pipeline [78]. After QC, normalization and cell-type annotation, we were left with a data-matrix defined over 13905 protein coding genes and 11059 breast epithelial cells, of which 2434 derived from the pretreatment stage, 8321 from the posttreatment stage and 304 from the metastatic sample.
*6) Serial sampling of CLL patient:* This CITE-Seq longitudinal dataset is available from GEO GSE152469, and describes the transcriptional evolution of leukemic and immune cells of a CLL patient under ibrutinib treatment [72]. We focused only on the scRNA-Seq component of the data, using an already Seurat-normalized data object downloadable from the CellResDB resource (https://cellknowledge.com.cn/cellresponse/index.html) [109]. Briefly, this normalized dataset consists of 9230 cells and 15965 genes, encompassing 8692 B-cells, 24 CD4+ T-cells, 428 CD8+ T-cells, 42 Hematopoietic Stem cells and 44 NK-cells. We only considered the B-cell population which was subdivided further into 4048 pretreatment and 4644 posttreatment cells.

We also analyzed a large sci-RNA-Seq3 dataset (Mouse organogenesis cell atlas-MOCA) comprising around 2 million cells [69], in order to further validate that CCAT is a measure of developmental potential [66, 67]:

*7) MOCA mouse organogenesis*: We downloaded the cell-type annotation and count data from GEO GSE119945 and processed the data with Seurat pipeline. We focused on 1,782,179 cells encompassing 38 cell-types with data from 5 stages in embryonic development: E9.5 (n=141,889), E10.5 (n=335,247), E11.5 (n=541,172), E12.5 (n=396,853) and E13.5 (n=367,018).

In order to refine our colon and stomach TF-differentiation networks, we analyzed scRNA-seq data from the Tabula Sapiens [58]:

*8) Tabula Sapiens colon and stomach atlases:* The normalized scRNA-Seq datasets are available for download from https://tabula-sapiens.sf.czbiohub.org. Specifically, we downloaded the *h5ad files for colon and stomach, containing the counts and normalized data and cell-type annotations. In the case of stomach and colon, the resulting expression matrices were defined for 33,064 and 13,680 normal cells, respectively. In the case of stomach, these 33,064 cells were 550 endothelial, 1601 epithelial, 6545 fibroblasts, 24245 immune-cells with the rest being smooth muscle (n=27), enterocyte-endocrine (n=25) and interstitial Cajal cell (n=71). In the case of colon, the 13,680 cells were distributed as fibroblasts (n=953), immune cells (n=6454), endothelial (n=53) and epithelial (n=6220).

To study dynamics of fetal gene expression in cancer progression we studied the atlas of human fetal gene expression (DESCARTES) [85], supplementing this with a fetal lung atlas [110].

*9) DESCARTES human fetal atlas:* This is a single-cell combinatorial indexing for gene expression (sci-RNAseq3) dataset of 121 human fetal samples ranging from 72 to 129 days in estimated postconceptual age, representing 15 organs and around 4 million cells. We downloaded the processed files *DE_gene_77_main_cell_type.csv* and *DE_gene_by_organ.csv* from https://descartes.brotmanbaty.org/bbi/human-gene-expression-during-development/ which contain differential expressed gene lists. Briefly, we constructed fetal gene expression lists for various cell-types in different organs using the within organ as well as between organ DEG-lists, requiring FDR<0.05 and ideally fold-changes > 2 or 1.5. In the case of lung, this yielded a 65-gene fetal squamous, a 438-gene fetal cilia and a 31-gene fetal alveolar signature. Because the number of fetal lung basal cells in DESCARTES was too small, the fetal basal signature was derived from the fetal lung atlas [110] by performing DEG-analysis (Wilcoxon tests) of basal cells to all other cell-types. This resulted in a 65-gene fetal basal signature. From DESCARTES we also derived fetal gene expression signatures for stomach goblet (n=39 genes) and chief (n=96) cells, as well as colon epithelial cells (n=63).

### Bulk gene expression datasets

We analyzed two bulk lung tissue gene expression datasets. One was generated with Agilent arrays and profiled a total of 98 samples encompassing every stage in lung squamous cell carcinoma progression [93]: normal (n=22), hyperplasia (n=13), metaplasia (n=13), dysplasia (n=27), lung carcinoma in-situ (n=9) and invasive cancer (n=14). Intra-sample normalized data was downloaded from GEO GSE33479. Inter-sample normalization was performed using quantile normalization as implemented in limma [111]. The other dataset derives from the TCGA LSCC [94], encompassing 45 normal-adjacent samples and 473 lung squamous cell carcinomas. We used the log-normalized data as described by us previously [49, 112].

### Construction of TF differentiation networks

The procedure for constructing a tissue-specific TF-differentiation network is described in detail in our previous publications [44, 49]. Briefly, the procedure uses a machine-learning algorithm called SCIRA to derive, for any given tissue-type, a number of tissue-specific TFs and associated TF-regulons, using the large GTEX v1 dataset encompassing 8555 samples and 29 tissue-types [46]. We focused on the older v1 version, because it is already appropriately powered [44] and because SCIRA is sensitive to potential batch effects present in the newer versions of GTEX. Moreover, the TF-differentiation networks that we have previously derived for lung and esophagus tissue using GTEXv1 have been extensively validated in previous work [44, 45], demonstrating the quality of the inference. In SCIRA, the TFs are identified by overexpression analysis comparing the given tissue-type of interest to all other tissue-types. To avoid confounding by immune cell infiltration, the overexpression analysis is carried out again by comparing the given tissue to blood, which we found to be a very effective procedure to remove TFs specific to resident immune-cells in cases where tissue-types have significant immune-cell infiltration [44]. To find the regulons of each TF, SCIRA implements a greedy partial correlation strategy, which identifies genes that are highly correlated with the TF’s expression level. As shown by us previously in the case of lung and esophagus, these gene regulons are enriched for direct binding targets of the TF, but they also contain indirect downstream targets that faithfully represent upstream binding activity [44]. In this work, we used SCIRA to derive TF differentiation networks for colon and stomach tissue, which resulted in 56 colon-specific and 52 stomach-specific TFs. The average regulon size in the case of stomach-specific TFs was 38 genes (range: 11-156), whereas in the case of colon-specific TFs, the average regulon size was 34 genes (range: 10-112).

### Estimation of transcription factor differentiation activity (TDA)

The differentiation activity of a given TF (here called TDA) is obtained as the t-statistic of a linear regression of a sample’s expression profile, be it a bulk RNA-Seq sample or a single-cell (scRNA-Seq), against the binding regulon profile of the TF, where positive and negative targets are encoded as +1 and -1, respectively, and with all non-targets set to 0. This estimation procedure has been extensively validated by us [44, 45].

### Refinement and validation of TF differentiation networks

Whilst SCIRA successfully identifies tissue-specific TFs, it still leaves open the possibility that a fraction of the identified TFs are specific to other cell-types within the tissue, such as endothelial cells or fibroblasts, specially if the tissue under consideration has a relatively high number of these. Thus, here we refined the colon and stomach-specific networks by removing TFs that we could not validate in the large scRNA-Seq Tabula Sapiens (TS) Atlas dataset [58]. In more detail, we used the colon-specific TF-regulons to estimate the TF’s TDA levels in the epithelial, immune, endothelial and fibroblast cells from the normal colon dataset of TS (n=13,680 cells). TFs’s were then called ‘colon-epithelial-specific’ if their TDA was significantly higher in epithelial cells compared to endothelial, immune and fibroblasts. A similar procedure was carried out for the gastric/stomach specific TFs using 33,064 normal cells from the stomach TS set. After this filtering, we were left with 18 colon-specific TFs and 47 gastric-specific TFs.

To further validate the TF-regulons we tested for enrichment of the regulon genes for direct TF-binding targets using the ChIP-Seq atlas [113] (http://chip-atlas.org), or if ChIP-Seq data was not available, against other high-quality integrated regulon databases such as DOROTHEA [60]. We stress that although the DOROTHEA regulons are very general regulons and not tailored for estimating TF-differentiation activity, that nevertheless some overlap and enrichment should be expected, specially for TFs that do not display functional pleiotropy. In the case of ChIP-Seq data, we checked if the binding intensity of the predicted regulon genes is higher than for non-regulon genes using a one-tailed Wilcoxon rank sum test. The binding intensity of each gene was averaged over samples, restricting to TFs for which ChIP-Seq profiles have been generated in at least 5 different samples, excluding embryonic samples, hESCs and predictions from the STRING database. In the case of DOROTHEA, we computed a one-tailed Binomial test P-value to assess if the overlapping number of genes between our TF-regulons and the DOROTHEA ones is statistically significant.

### The PLASTIC (Predictive Landscape Analysis of Single-cell Transcriptomes for Identifying Cancer) framework

#### Construction of the Waddington attractor landscape

PLASTIC is aimed for scRNA-Seq/snRNA-Seq studies that have profiled normal tissue, histologically normal-tissue at cancer-risk, and ideally also cancer-tissue itself. Normal tissue at cancer-risk could include diverse types of precancerous lesions, for instance, metaplasia, dysplasia, polyps or even normal-tissue exposed to a major cancer risk factor (e.g. smoking). Once the tissue-type is fixed, it first uses the tissue-specific TF differentiation network, as derived earlier, to estimate TDA-levels for the tissue-specific TFs in each single-cell. This results in a TDA-matrix defined over a given number of tissue and cell-type specific TFs (usually epithelial specific as we are interested in epithelial cancer development) and cells. From this TDA-matrix we then construct a diffusion map [61, 114] in order to identify potential attractor states and the diffusion paths connecting them. Focusing on the first two diffusion components (DC1/2) is usually sufficient to obtain a reasonable representation of the cellular states involved. This two-dimensional DC-space is defined within the TDA-space itself and can be used to representing the 2D projection of a Waddington-like landscape. To define the elevation in this landscape, we use CCAT [66], an ultra-fast approximation to diffusion entropy [62], which can be viewed as measuring general cellular plasticity. This is based on the observation that it correlates very well with developmental potential across multiple species [67], with differentiation potential across multiple different lineages (both single-cell and bulk) [62, 64, 66], with clinical outcome in cancer [115], cell-fate commitment within unipotent lineages [45], as well as drug resistance [62]. Briefly, CCAT is defined by the Pearson Correlation Coefficient (PCC), i.e. 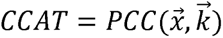, between a single cell’s genome-wide RNA-Seq profile 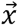 (normalized and log-transformed) and the connectivity (i.e. degree or number of neighbors) profile, 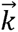, of the corresponding proteins as determined by a highly curated protein-protein interaction (PPI) network [116, 117] from Pathway Commons [118, 119]. This simple formula can be derived as an approximation to the Diffusion/Signaling Entropy Rate (*SR*), also called SCENT [62], which is given by the formula

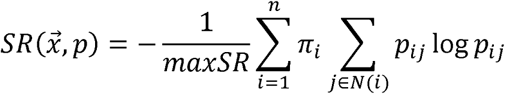

where are the entries of a stochastic matrix, and _π_ is the invariant measure, satisfying π*P=*π and the normalization constraint π^T^**1***=*1. The stochastic matrix is given by the formula

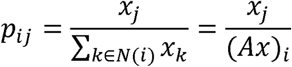

where *N(i)* denotes the neighbors of protein *i,* and where *A* is the adjacency matrix of the PPI network (*A_ij_=1* if *i* and *j* are connected, 0 otherwise, and with *A_ii_=*0). The reason why CCAT measures plasticity is that cells displaying higher phenotypic plasticity tend to overexpress network hubs, with many of these network hubs encoding ribosomal proteins [62], a result we have validated across over 2 million cells and 28 scRNA-Seq studies [66]. Of note, the association between ribosomal gene expression and differentiation potential is observed across different species and is independent of cell-proliferation [81, 82].

Having estimated the cellular plasticity of each cell (and hence for each attractor state), an approximation to Waddington’s landscape is constructed as follows. First, one must identify the cells making up the attractors, which is accomplished using ordinary techniques such as cell-type annotation following dimensional reduction and/or using information about the sample (i.e. was it derived from cancer-tissue, or metaplasia). Using the diffusion components DC1 and DC2 as well as the estimated CCAT values, we can thus obtain estimates of the positions of the attractors in a 3D landscape. Since attractors are normally depicted as minima/basins, we use the label *m.* Hence, for each minimum *m* we have a position (x_m_, y_m_, z_m_) with *(x,y)* representing DC1/2 and *z* representing CCAT. Next, we need to estimate the shallowness (standard deviation) of the attractors, which is done based on the diffusion map values. Let denote the standard deviation of the attractor *m.* To generate the Waddington landscape (i.e. surface) we then define the following function z=*W(x,y)* where

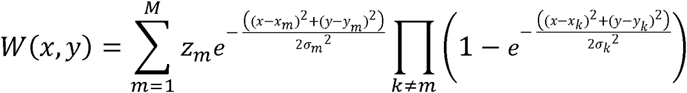

and where *M* is the number of minima/attractors. This function satisfies all the requirements: (i) it displays a local minimum at each attractor state (x_m_, y_m_), (ii) the height of each attractor is proportional toz*_m_*(unless attractors are very close to each other, the product term above is approximately equal to 1), (iii) it ensures smoothness of the surface.

#### Calculation of the Transcription Factor Inactivation and Activation Loads (TFIL/TFAL)

To quantify epigenetic reprogramming (i.e. dedifferentiation or transdifferentiation), we use the following TFIL/TFAL metrics. For each cell, the TFIL is computed by counting the number of tissue-specific TFs that are inactivated in that cell compared to the normal cells. By inactivation we mean that the TDA value of a given TF is significantly lower in the given cell compared to a null distribution defined by the TDA values of the normal cells. Because TDA-values are non-sparse (they are t-statistics), we approximate the null distribution by a Gaussian of a given mean and standard deviation. For each cell, we then compute the z-score deviation from this null, and if the nominal P-value < 0.05, we declare the TF as inactivated in that cell. We use a relaxed threshold, but more stringent thresholds are acceptable, as the aim is to simply rank cells by their TFIL values. The TFAL is computed in an analogous fashion, by calculating the number of TFs that display increased TDA in a given cell compared to the normal cells. However, for TFAL, the set of TFs to use would be different compared to those used in TFIL, as the aim of computing TFAL is to estimate transdifferentiation into another cell-type. For instance, in Barret’s Esophagus we would use esophagus squamous epithelial specific TFs to estimate TFIL, whereas for TFAL we would consider colon-epithelial specific TFs, since Barret’s is an intestinal form of metaplasia.

#### Calculation of the cancer-risk score (CRS)

To estimate the cancer-risk score of a cell, we use a Kernel function to describe the similarity of a cell (usually a precancerous cell) to each cancer cell. This similarity is computed in TDA-space. Here we compute the CRS using a simple Pearson Correlation between the cell’s TDA profile and that of each cancer-cell. To arrive at one CRS-value for each precancerous cell, we take the average of the PCC-values over all cancer-cells. We point out that the CRS and TFIL values calculated for each precancerous cell are independent measures, because the TFIL is estimated by comparison to the normal state, whereas the cancer-risk score reflects the similarity to the cancer cells. Thus, a positive association between TFIL and the cancer-risk score is non-trivial and would indicate that precancerous cells with a higher TFIL are more similar to cancer cells.

#### Calculation of cell-cycle scores

To identify single cells in either the G1-S or G2-M phases of the cell-cycle we followed Tirosh et al [120]. A given normalized scRNA-Seq data matrix is first z-score normalized for all genes defining the G1-S or G2-M phase signatures. A cycling score for each phase and each cell is then obtained as the average z-score over all genes present in each signature.

#### PPI network

We downloaded the integrated human PPI network from Pathway Commons (www.pathwaycommons.org) encompassing, 6692 pathways, 3,579,336 interactions curated from 23 databases [118, 119]. Specifically, we used the PC2v12 version. The network was further filtered by using main cellular localization data from the HPRD, to remove interactions between proteins that do not map to neighboring signaling domains, as described by us previously [62]. The end result is a 12,649 protein network that is connected. The network is sparse with 0.5% interactions (i.e. 928,182 PPIs).

### Differential gene expression analyses and lung basal CRS signature score

Differential gene expression analyses between high and low cancer risk states were performed using Wilcoxon rank sum tests. In the case of the lung smoking study, we identified non-cycling cells as cells with negative G1-M and G2-S cell-cycle scores. Non-cycling cells with TFIL >1 were then compared to non-cycling cells with TFIL=0 and differentially expressed genes identified using Benjamini-Hochberg FDR<0.05. The top 300 upregulated and 300 downregulated genes were used for GSEA as well as for defining a cancer-risk signature and score (CRS). For independent data, the CRS-score was obtained by performing a Pearson correlation between the dichotomized signature (1 for upregulated genes, -1 for downregulated genes) and the expression profile of the sample. This was done after z-score normalizing the independent mRNA expression dataset.

### Gene Set Over-enrichment Analysis

As a database of biological terms we used MSigDB_2023v2, which contains 34,555 terms, including 50 cancer-hallmark gene sets, 301 positional gene sets, 7233 curated gene sets, 3713 regulatory target gene sets, 1007 cancer-oriented module gene sets, 16008 gene-ontology sets, 189 oncogenic perturbation signatures, 5219 immunological perturbation/state signatures and 830 cell-type specific marker sets. To test for enrichment, we used a one-tailed Fisher-test, as implemented in the ebGSEA R-package [86].

## Code Availability

PLASTIC is an R-package that is freely available from https://github.com/aet21/plastic.

## Acknowledgements

This project was funded by the National Natural Science Foundation of China grants 32170652 and 32370699, as well as a Research Fellowship for International Scientists (RFIS, W2431024).

## Author contributions

Study was conceived by AET. Statistical and bioinformatic analyses were performed by JTHK and AET. Manuscript was written by AET with contributions from JTHK.

## Ethics approval

Not applicable as we only analyzed publicly available data.

